# Evaluation of NGS-based approaches for SARS-CoV-2 whole genome characterisation

**DOI:** 10.1101/2020.07.14.201947

**Authors:** Caroline Charre, Christophe Ginevra, Marina Sabatier, Hadrien Regue, Grégory Destras, Solenne Brun, Gwendolyne Burfin, Caroline Scholtes, Florence Morfin, Martine Valette, Bruno Lina, Antonin Bal, Laurence Josset

## Abstract

Since the beginning of the COVID-19 outbreak, SARS-CoV-2 whole-genome sequencing (WGS) has been performed at unprecedented rate worldwide with the use of very diverse Next Generation Sequencing (NGS) methods. Herein, we compare the performance of four NGS-based approaches for SARS-CoV-2 WGS. Twenty four clinical respiratory samples with a large scale of Ct values (from 10.7 to 33.9) were sequenced with four methods. Three used Illumina sequencing: an in-house metagenomic NGS (mNGS) protocol and two newly commercialized kits including a hybridization capture method developed by Illumina (DNA Prep with Enrichment kit and Respiratory Virus Oligo Panel, RVOP) and an amplicon sequencing method developed by Paragon Genomics (CleanPlex SARS-CoV-2 kit). We also evaluated the widely used amplicon sequencing protocol developed by ARTIC Network and combined with Oxford Nanopore Technologies (ONT) sequencing. All four methods yielded near-complete genomes (>99%) for high viral loads samples, with mNGS and RVOP producing the most complete genomes. For mid viral loads, 2/8 and 1/8 genomes were incomplete (<99%) with mNGS and both CleanPlex and RVOP, respectively. For low viral loads (Ct ≥25), amplicon-based enrichment methods were the most sensitive techniques yielding complete genomes for 7/8 samples. All methods were highly concordant in terms of identity in complete consensus sequence. Just one mismatch in two samples was observed in CleanPlex *vs* the other methods, due to the dedicated bioinformatics pipeline setting a high threshold to call SNP compared to reference sequence. Importantly, all methods correctly identified a newly observed 34-nt deletion in ORF6 but required specific bioinformatic validation for RVOP. Finally, as a major warning for targeted techniques, a default of coverage in any given region of the genome should alert to a potential rearrangement or a SNP in primer annealing or probe-hybridizing regions and would require regular updates of the technique according to SARS-CoV-2 evolution.

## Introduction

A novel human betacoronavirus, Severe Acute Respiratory Syndrome Coronavirus 2 (SARS-CoV-2), emerged in China in December 2019, rapidly spreading worldwide and resulting in the coronavirus disease 2019 (COVID-19) pandemic (1). Whole-genome sequencing (WGS) of SARS-CoV-2 has played a major role since the onset of the pandemic. Notably WGS has contributed to design specific RT-PCRs (2), antiviral strategies (3), and vaccine candidates (4). SARS-CoV-2 WGS also allowed to explore lineage transmission and might be useful to assess the effectiveness of intervention measures (5–8). Furthermore SARS-CoV-2 genomic surveillance allowed the characterization of some mutations for which the impact on virulence and transmissibility needs to be confirmed (9–11). SARS-CoV-2 WGS is performed at an unprecedented rate worldwide with the use of very diverse Next Generation Sequencing (NGS) methods. Viral metagenomic NGS (mNGS) enabled early sequencing of the SARS-CoV-2 genome (1) and is the method routinely used in the French National Reference Centre for Respiratory Viruses (NRC, Lyon, France). However, this method lacks sensitivity and cannot produce whole genomes for low viral load samples, except using a very high depth of sequencing (10). Targeted methods such as amplicon- or capture-based enrichments are promising candidates to overcome this issue. A formal comparative study of these different approaches has not been performed so far. Herein, we aimed to assess performance of four NGS-based approaches for SARS-CoV-2 WGS. Three used Illumina sequencing: an in-house metagenomic NGS (mNGS) protocol and two newly commercialized kits including a hybridization capture method developed by Illumina and an amplicon sequencing method developed by Paragon Genomics. We also assessed a widely used amplicon sequencing protocol developed by the ARTIC network and combined with Oxford Nanopore Technologies (ONT) sequencing (7,12). The present evaluation covers experimental process as well as the associated bioinformatic solutions recommended for each technique. Importantly, this evaluation included two samples with a newly observed 34-nt deletion in ORF6 [Queromes et al, in preparation] in order to assess the capacity to detect large modification in SARS-CoV-2 genome.

## Methods

### Sample selection

A total of 24 clinical samples (nasopharyngeal swab) with a broad range of representative SARS-CoV-2 cycle threshold (Ct) values (from 10.7 to 33.0) were selected for sequencing. We defined 3 groups of samples: a low Ct values group (Ct<20), a medium Ct values group (20<Ct<25) and high Ct values group (Ct≥25). Each sample was tested with 2 real time reverse-transcriptase polymerase chain reactions (RT-PCR) targeting distinct RdRp gene regions (Institut Pasteur assay, Paris, France (13)). Total nucleic acid extraction was performed with easymag platform (bioMérieux, Lyon, France). Samples used in this study were collected as part of approved ongoing surveillance conducted by the NRC at the Hospices Civils de Lyon. The investigations were carried out in accordance with the General Data Protection Regulation (Regulation (EU) 2016/679 and Directive 95/46/EC) and the French data protection law (Law 78–17 on 06/01/1978 and Décret 2019–536 on 29/05/2019).

### Sequencing approaches and technologies

### Illumina sequencing

#### Viral metagenomics (mNGS)

DNase treatment (Life Technologies, Carlsbad, CA, USA) was performed after nucleic acid extraction in order to increase sensitivity of RNA virus detection and overcome human contamination (14). Nucleic acids were randomly amplified using the WTA2 kit (WTA2, Sigma-Aldrich, Darmstadt, Germany) and libraries were prepared using the Illumina Nextera XT kit (Illumina, San Diego, CA, USA). This in-house approach is routinely used to perform WGS SARS-CoV-2 genomic surveillance at the NRC (10,15) for samples with a Ct value <20.

#### Hybrid capture-based target enrichment

Hybrid capture-based approach was performed using the Illumina DNA Prep with Enrichment kit and Illumina Respiratory Virus Oligo Panel (RVOP) V1.0. Briefly, double stranded cDNA synthesis was performed from DNase pretreated total nucleic acid extracts using the NEBNext^®^ Ultra™ II RNA First Strand Synthesis Module and NEBNext^®^ Ultra™ II Non-Directional RNA Second Strand kit (New England Biolabs, MA, USA). The enrichment was based on a hybridization step with a respiratory viruses biotinylated adjacent oligoprobes panel (16) recently expanded to include SARS-CoV-2.

#### Amplicon-based target enrichment

We implemented the CleanPlex SARS-CoV-2 panel (Paragon Genomics, Inc, Hayward, CA, USA) protocol for target enrichment and library preparation (17). In short, from reverse-transcribed RNA, multiplex PCR reactions were performed using 343 pairs of primers separated into two pools covering the entire genome of SARS-CoV-2 ranging from 116bp to 196bp, with a median size of 149 bp. Illumina indexes were introduced by PCR.

For these three approaches the prepared libraries were sequenced on an Illumina NextSeq™ 550 with mid-output 2×150 bp (mNGS and CleanPlex) or 2×75 bp (RVOP) flow cells.

### ONT sequencing using amplicon-based target enrichment

We tested a multiplexed PCR amplicon approach implementing the ARTIC Network nCoV-2019 sequencing protocol slightly modified by ONT (Oxford Nanopore Technologies, Oxford, UK) for better performance. The multiplex PCR primers set v3 was used to span the whole genome (12). Briefly, synthesized cDNA was used as template and tiling 400nt-amplicons with 20 base pairs overlaps (not including primers) were generated using two pools of primers for 35 cycles. Samples were multiplexed by using the native barcode kits from ONT (EXP-NBD104 and EXP-NBD114). The library was prepared using the SQK-LSK109 kit and then sequenced on a FLO-MIN106 (R9.4.1) flow cell, multiplexing 24 samples per run.

### Sequencing data analysis

mNGS data were analysed with an in-house pipeline. Briefly, low quality and human reads were filtered out (dehosting) and remaining reads were aligned to the SARS-CoV-2 reference genome (isolate Wuhan-Hu-1, EPI_ISL_402125) using the BWA-MEM (v0.7.15-r1140) algorithm. Consensus sequences were generated through a simple majority rule using custom perl script. These sequences were used as the patients’ own mapping reference for further realignment of the reads. Final consensus sequence was called at 10x using no-clip alignment. Visual inspection of read alignments was performed using Integrative Genomics Viewer (IGV) (18) around regions with large drop in coverage to detect and define potential deletions. For the CleanPlex SARS-CoV-2 amplicon approach, data analysis was performed according to the supplier’s recommendations with a pipeline developed by Sophia Genetics (V1) (17). Sequencing data from the RVOP capture approach were parsed using the Illumina Dynamic Read Analysis for GENomics (DRAGEN; v3.5.13; Illumina) Bio-IT Platform. Finally, ONT sequencing data were analysed by implementing the recommended bioinformatics developed by ARTIC and available online (v1.1.0) (19). Briefly, basecalling and demultiplexing were performed using Guppy high accuracy models (v3.5.2). Local alignment was performed using Minimap2 (v 2.17) and variant calling with nanopolish (v0.13.2). Five parameters were specifically evaluated and compared: coverage metrics such as depth and breadth of coverage at 10x, number of SNPs between methods after pairwise alignment of consensus sequences, and detection of a particular 34nt-deletion detected with our in-house reference mNGS method. The proportion (%) of genome coverage are presented as medians with interquartile ranges [IQR] and compared using the non-parametric Friedman test.

## Results

### Sensitivity assessment of four SARS-CoV-2 sequencing approaches

Four widely used NGS methods were evaluated: an unbiased mNGS approach, and three targeted approaches based on hybridization capture (RVOP) or amplicon sequencing (CleanPlex and ARTIC-ONT). For high viral loads (Ct <20), the 4 sequencing approaches yielded almost-complete genome (> 99 % covered at 10x) for all samples. A significant difference in genome coverage distribution (p<0.001) was found; the highest median of coverage was for RVOP and mNGS (Table 1). For mid Ct samples, one sample sequenced with RVOP and two samples sequenced with mNGS had a genome coverage below 99% (93.4% with RVOP, 72.6% and 56.5% with mNGS, respectively).

Among samples with low viral loads (Ct>25Ct), significant discrepancies in genome coverage were observed between methods (p=0.005); 8.8% median coverage in samples sequenced with mNGS, 92.0% for RVOP, and highest median coverage were obtained with amplicon sequencing (99.4%, CleanPlex; 99.6%, ARTIC-ONT).

While RVOP and mNGS enabled to generate the complete genome (maximum coverage was 100% for highest viral loads), amplicon-based target enrichment did not allow to cover SARS-CoV-2 genome ends, as expected by considering the design of ARTIC-ONT and CleanPlex protocols. The highest coverage was 99.6% and 99.7% for ARTIC-ONT and CleanPlex, respectively.

Irrespective of Ct value, RVOP provided an even depth of coverage throughout the genome, while uneven depth of coverage was obtained with mNGS and marked drops in sequencing depth were observed for CleanPlex (Figure 1). We did not interpret the evenness of depth of coverage for ARTIC-ONT as the dedicated pipeline limits the depth at 400x. These drops corresponded to amplicons that were poorly amplified: from 5,159 to 5,199 (drop # 1), from 14,430 to 14,505 (drop # 2), from 19,337 to 19,399 (drop # 3), and from 22,641 to 22,715 (drop # 4; Figure 1). Regarding low and mid Ct samples, 1, 3, and 8 had a coverage <10x at drops # 2, #1, and # 4, respectively. Importantly none of these regions involved crucial positions used to define the Nextstrain major clades (20).

**Figure 1.**
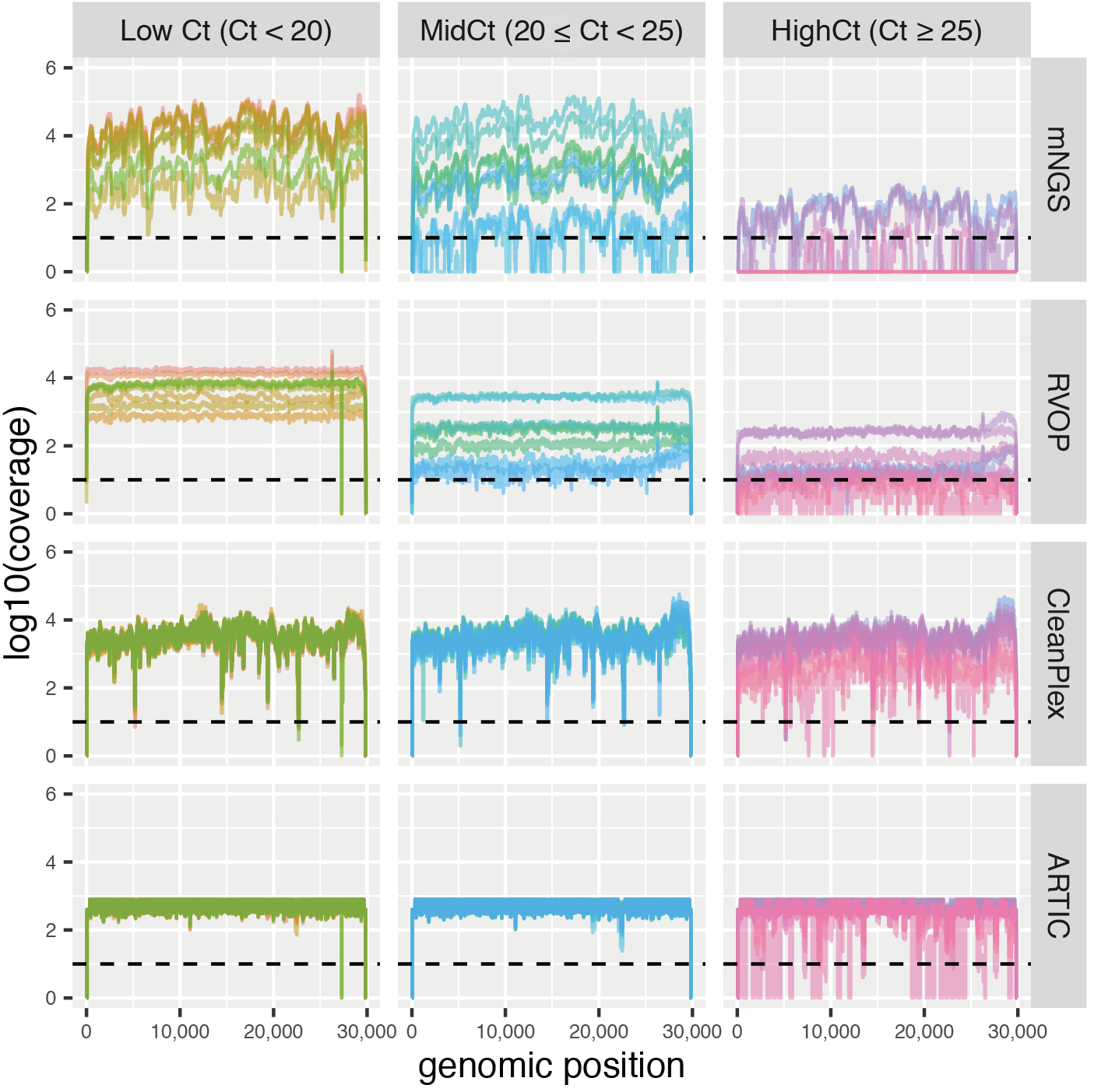
Plots of coverage according to evaluated methods and Cycle threshold (Ct) values groups. Using an R script, these plots were constructed via ggplot2 from depth files generated by bedtools from output aligned bam files of each specific-method pipeline. Of note, ARTIC-ONT pipeline limits the depth at 400x, the real depth is not represented herein.

### Comparison of complete genome consensus sequence

Almost-complete genomes generated with the 4 methods were further compared to define accuracy of each approach (Figure 2). Of note, 9, 7, and 3 samples sequenced, respectively, with mNGS, RVOP and CleanPlex were excluded of the analysis due to a coverage <99% at 10x, *vs* 1 sample for ARTIC-ONT protocol. For all other samples, all consensus sequences were strictly identical, expect for 2 samples for which CleanPlex led to one mismatch compared to the other methods (sample # 1, position 280086; sample # 20, position 11083). By analysing VCF files, these 2 positions corresponded to SNPs (G280086T, G11083T) with variant frequency <70% which is the threshold set for calling a SNP in Sophia Genetics pipeline while other pipelines set this threshold at 50%.

**Figure 2.**
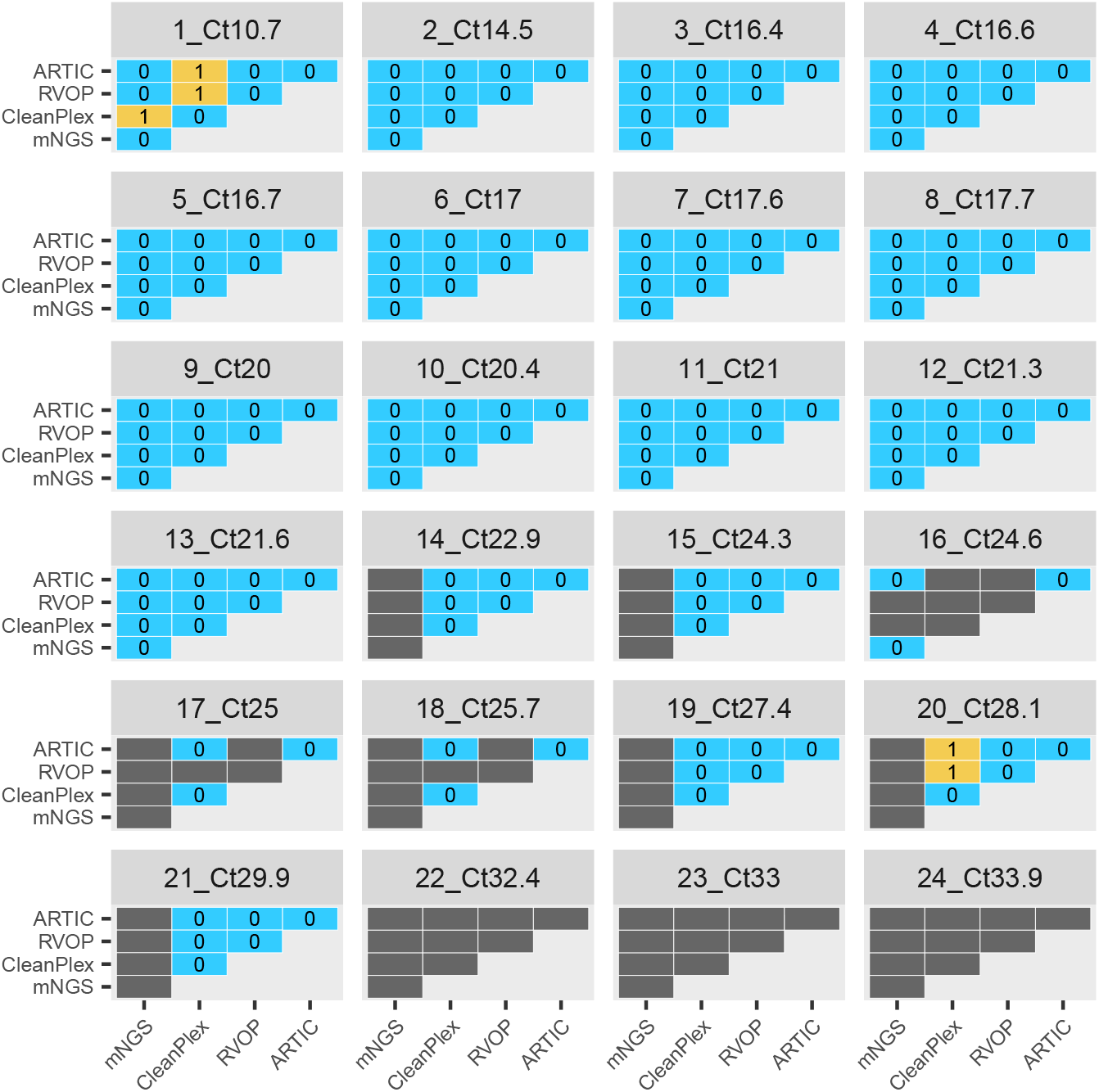
Mismatch count between consensus sequences generated by each method compared 2 by 2 for each sample. These matrices were generated only from consensus with determined bases for more than 99% of the genome. If one sequence of the two had more than 1% of undetermined bases (N), comparison was not assessed, grey tiles. Blue tiles correspond to perfect identity and orange tile correspond to mismatches, the number of mismatches is indicated inside the tile. Matrices were generated with an R script using Decipher (alignment), ape (distance matrices) and ggplot2 (charts) libraries. Of note, undetermined bases and deletions were not considered in the calculation of mismatches.

### Performance of the methods for the detection of a large deletion in SARS-CoV-2 genome

A 34 nt-deletion in the ORF6 at loci 27266 was detected in 2 samples (samples # 7 and # 8) by all the evaluated approaches both in alignment files and consensus sequences. For RVOP, it is noteworthy that DRAGEN output alignment files showed an absence of coverage in the region but there were no spanning reads to visualise the breakpoint of the deletion. However, DRAGEN output variant calling files clearly highlighted the 34 nt-deletion as well as the final consensus sequence for both samples. Using Minimap2 as aligner, the deletion was clearly identified with reads spanning the breakpoint of the deletion on the reference genome (Supplementary figure 1).

## Discussion

We present herein the evaluation of 4 representative NGS-based approaches for SARS-CoV-2 whole genome characterisation. Unbiased mNGS is the most appropriate method to identify novel emerging pathogens such as SARS-CoV-2 (1,21) but did not produce all WGS for mid and low Ct values samples in the present study. In contrast, the three targeted methods evaluated herein exhibited higher sensitivity compared to mNGS by reaching a high proportion of the genome covered. In particular, amplicon-based target enrichment using CleanPlex or ARTIC-ONT was the most sensitive approach and allowed WGS of samples with Ct up to 33.9. Although not reported herein, amplicon-based target enrichment can be impacted by SNP or indels located within primer-annealing regions even if their tiling amplicon designs (instead of adjacent amplicon designs) aims to reduce the impact of such modifications. With SARS-CoV-2 evolving, primers for amplicon-based target enrichment need to be constantly updated, as it has been done for ARTIC-ONT (v3) (22). Such updates will be important for CleanPlex as we have observed coverage dropout issues in 4 regions of the genome irrespective of Ct value, suggesting potential mismatches in primer-annealing regions that may lead to variation in the optimal annealing temperature and thus to a decrease in the amplification efficiency.

All 4 methods were highly concordant in terms of identity in complete consensus sequence. Just one mismatch in two samples was observed in CleanPlex *vs* the other methods, due to the dedicated pipeline setting very a high threshold to call SNP compared to reference genome. Considering the very low evolution rate of this virus that can be explained by the editing function of its polymerase, just one mutation can have high repercussions in terms of evolutionary assessment.

Furthermore, a novel 34nt-deletion in the ORF6 previously observed with our in-house mNGS method, was detected by the two evaluated tiling amplicon sequencing methods (ARTIC and CleanPlex). However larger deletions spanning two primer annealing regions may arise and would be difficult to define using amplicon-based target enrichment methods. Missing regions with no coverage should be carefully investigated using other methods. The 34nt-deletion was also detected using RVOP despite an adjacent oligoprobes design. As the oligoprobe set is larger than the primer panel used in tiling amplicons protocols, a more comprehensive profiling of regions is obtained independently of genomic rearrangement. Moreover, a major advantage of this capture protocol, is the enrichment of other respiratory viruses that enables to detect coinfections (16).

The present study does have several limitations. First, the 4 methods were tested on a limited number of European samples, all affiliated to the lineage B1, according to the Pangolin classification (23); the present evaluation should be confirmed in larger studies including other lineages in order to be more representative of the global ecology of this emerging virus. As SARS-CoV-2 sequencing is an expanding market and other methods has been published (24–26) or are probably under development, the present study is not exhaustive. However, the approaches selected herein are representative of the methods mostly used during pandemics and are easy to implement in a diagnostic laboratory. As mentioned above, dedicated bioinformatic processes have been implemented as recommended by suppliers. Except for RVOP for which we compared the aligner using in DRAGEN and minimap2, we chose not to perform a comparison of the bioinformatic tools. In this context, further bioinformatic evaluations are needed especially concerning parameters such as the variant frequency threshold for calling SNP compared to reference sequence, as well as the minimum depth for calling consensus. Additionally, *de novo* assembly should be evaluated as it may provide better assessment of rearrangements such as large deletions, insertions and duplications that can be missed by classical alignment analyses, but this method is time-consuming and may not be appropriate for genomes with low coverage. More broadly, a repeatability test is mandatory to comfort results obtained from the present evaluation. Finally, cost effectiveness and turnaround time were not assessed in this study considering that many parameters such as laboratory equipment and prices negotiated with suppliers may affect results.

Beyond these limitations, the data presented herein are useful for clinical and research teams who want to implement SARS-CoV-2 sequencing and chose the most suitable protocol according to the application. To summarise, mNGS remains the gold standard for samples with high viral load to obtain a maximum of information without any bias. For low and mid Ct values, RVOP leads to very high coverage as well, enabling genome end sequencing, contrary to amplicon methods. For higher Ct values, amplicon-based enrichment are a very interesting alternative, in particular ARTIC-ONT protocol that did not show any dropout issues in this present evaluation. However, as a reminder, default of coverage in any given region of the genome should alert to a potential rearrangement or an SNP in primer annealing or probe-hybridizing regions and require regular updates according to SARS-CoV-2 evolution.

## Acknowledgements

We would like to thank all the patients, laboratory technicians and clinicians who contributed to this investigation. We are also grateful to Philip Robinson (DRCI, Hospices Civils de Lyon) for help in manuscript preparation.

## Statement of data availability

Data will be deposited on SRA database prior to publication.

## Legends

**Table 1.** Proportion of coverage throughout the SARS-CoV-2 genome at 10x according to evaluated methods. Cycles threshold (Ct) values were determined using the RdRp Institute Pasteur RT-PCR assay (IP4). Using an R script, these proportions (%) were calculated from depth files generated by bedtools from output aligned bam files of each specific-method pipeline. The percentage of genome coverage are presented as medians with interquartile ranges [IQR] and compared using the non-parametric Friedman test.

**Supplementary Figure 1.** Visualisation by Integrative Genomics Viewer (IGV) of the 34-nt deletion in the ORF6 highlighted from capture sequencing data of one of the two samples concerned (sample # 8). Plot of coverage from DRAGEN pipeline is represented in grey with conventional numbering systems for SARS-CoV-2 reference (MN908947.3). Depth at position 27079 is indicated into brackets. Details of output alignment files from DRAGEN did not show any reads (grey) overlapping the deletion contrary to those obtained using Minimap2 as aligner (black line within read).

