## Supplementary figures and images for "Evaluation of NGS-based approaches for SARS-CoV-2 whole genome characterisation"

### Supplementary figure

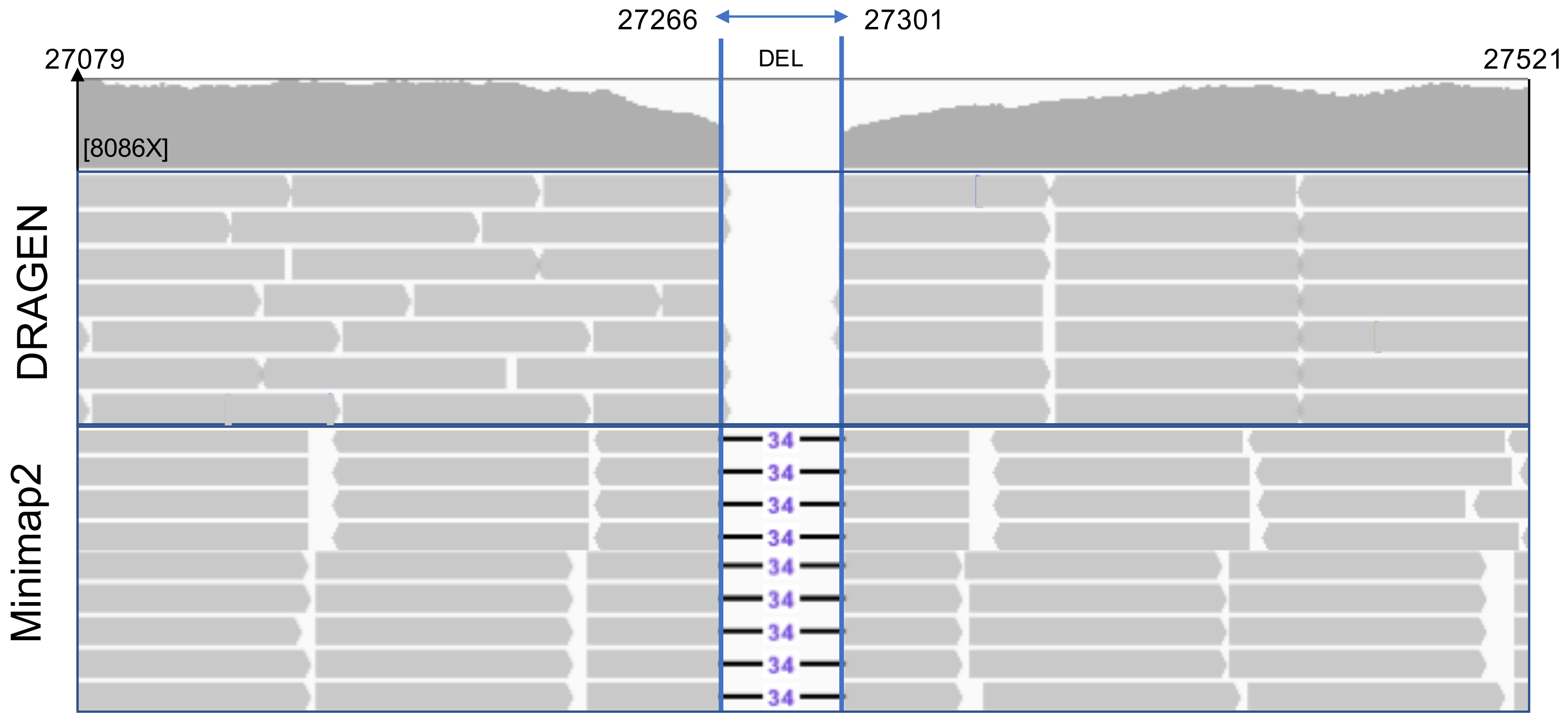
